# Comparative proteomic quantification reveals novel longevity regulators

**DOI:** 10.64898/2026.10.06.757189

**Authors:** Lang Ding, Hanfeng Lin, Cuizhen Huang, Viruthika Vallanadu, Kelly Ashley, Scarlett Rose Pitts, Ayse Sena Mutlu, Yue Zhou, Xing Ma, Jong Min Choi, Feng Jin, Jin Wang, Meng C. Wang

## Abstract

Proteomics is a powerful tool for profiling the protein landscape underlying cellular and physiological processes. Advances in mass spectrometry-based methods, such as tandem-mass-tags (TMT) isobaric labeling-based proteomics at both MS2 and MS3, label-free quantification (LFQ) with data-dependent-acquisition (DDA) or data-independent acquisition (DIA)-based proteomics, and stable isotope labeling with amino acid (SILAC) spike-in-assisted proteomics, enable quantitative proteome comparisons across biological conditions. These five quantitative proteomic methods offer distinct advantages in coverage, throughput, and quantitative reproducibility. Here, we treated them as orthogonal approaches, retaining protein candidates with significant, consistent-direction changes supported across multiple independently instrumented methods to identify high-confidence protein-level changes associated with longevity. Insulin/insulin-like growth factor signaling (IIS) is a highly conserved longevity regulatory pathway from *Caenorhabditis elegans* to humans. By quantitatively profiling IIS mutants in *C. elegans* and performing genetic screening, we obtained a systems-view of proteomic changes linked to longevity and identified new longevity regulators. This systematic cross-method comparison also serves as a valuable resource for guiding the selection of quantitative proteomics approaches.

## 1 | INTRODUCTION

As illustrated by the central dogma of molecular biology, genetic information flows from DNA to RNA to protein, highlighting proteins as the ultimate functional drivers of cellular and physiological processes (Ludvigsen & Honoré, 2018). Understanding the proteome is therefore crucial for bridging the gap between genotype and phenotype, providing direct insights into regulatory mechanisms underlying health and diseases (Sun et al., 2023). In addition, proteomic analysis reveals post-transcriptional and post-translational changes that are not necessarily captured by transcriptomic approaches (Takemon et al., 2021). However, quantitatively profiling the proteome has been technically more challenging than transcriptomics, but progress in mass spectrometry (MS)-based proteomics has led to significant advances in coverage, throughput, and quantification (Schmidt et al., 2011).

There are different strategies of quantitative bottom-up proteomics, including label-free quantification (LFQ), isobaric labeling quantification such as tandem mass tags (TMT), metabolic labeling quantification such as stable isotope labeling by amino acids (SILAC) (Fredens et al., 2011; Larance et al., 2011; Narayan et al., 2016; Thompson et al., 2003; Visscher et al., 2016; Walther et al., 2015). LFQ measures relative protein abundance by comparing the intensity of peptide signal across multiple independent mass spectrometry runs without employing isotopic or chemical labels (Cox et al., 2014). Isobaric labeling quantification chemically tags peptides from different samples with distinct mass labels, enabling multiplexed analysis in a single MS run and quantitation based on reporter-ion intensities from the MS2 or MS3 spectra (Mertins et al., 2018; Thompson et al., 2003; Ting et al., 2011). Metabolic labeling quantification incorporates stable isotope-labeled amino acids directly into proteins in living cells or organisms, providing an internally labeled standard for relative quantification between sample conditions (Larance et al., 2011; Ong et al., 2002). Few benchmark studies have compared these approaches. For example, it is reported that data-dependent acquisition (DDA)-LFQ coupled with the LFQ algorithm in MaxQuant can achieve accuracy comparable to SILAC (Cox et al., 2014). When comparing TMT-MS2 with TMT-synchronous precursor selection (SPS)-MS3 and SILAC, a previous study supports the use of TMT-MS2, as it offers much higher precision and deeper identification despite its lower accuracy (Hogrebe et al., 2018). While TMT offers the advantage of reducing mass spectrometer running time (Mertins et al., 2018; Thompson et al., 2003), potential experimental variations before TMT labeling cannot be excluded, along with interference and ratio distortion (Pappireddi et al., 2019). Applying SPS-MS3 would mitigate ratio distortion; however, it comes at the cost of reduced identification depth and quantification compared to the MS2 method (Ting et al., 2011). In addition, TMT and SILAC labeling are often associated with high reagent costs. In recent years, advances in bioinformatics have led to an explosion of data-independent acquisition (DIA) proteomic approaches, with improved software enabling DIA data analysis of complex MS2 spectrum (Bruderer et al., 2015; Demichev et al., 2020; Frohlich et al., 2022; Ludwig et al., 2018; MacLean et al., 2010; Sinitcyn et al., 2021; Tsou et al., 2015). To reach a depth comparable to TMT-MS2 and shorten mass spectrometer runtime, many strategies have been developed to optimize the DIA result, such as isolation window optimization (Ludwig et al., 2018), BoxCar DIA (Meier et al., 2018), and library-free DIA (Demichev et al., 2020). However, comprehensive cross-method comparisons using same biological samples remain limited. Meanwhile, Orbitrap mass spectrometers have rapidly evolved, and direct comparison of their performance could provide valuable insights into their applications in biological and biomedical research (He et al., 2023).

In the published literature, distinct proteomic approaches have been applied in different studies to identify differentially expressed (DE) protein candidates, and systematic comparisons are typically conducted using benchmarking controlled mixture. Thus, updated guidance for choosing quantitative proteomic workflows, with a focus on identifying DE protein candidates with sufficient coverage and confidence, is needed. We applied this strategy to characterize proteome changes in association with organism longevity. The insulin/insulin-like growth factor signaling (IIS) pathway is one of the most evolutionarily conserved longevity regulatory mechanisms, first discovered in *Caenorhabditis elegans* (*C. elegans*) and later shown to influence lifespan in various organisms, including human (Kenyon, 2010). In *C. elegans*, the loss-of-function mutation of *daf-2(lf)*, encoding the insulin/IGF-1 receptor, doubles the lifespan, which is fully suppressed by the loss-of-function mutation of *daf-16(lf)*, encoding the FoxO transcription factor in the IIS pathway (Kenyon et al., 1993; Kimura et al., 1997). A systemic understanding of the proteomic changes associated with this mechanism provide valuable information to the aging research community. Previous studies using SILAC based proteomics have profiled proteome dynamics across the lifespan of wild type (WT) worms as well as *daf-2(lf)* and *daf-16(lf)* mutants, identifying protein changes associated with aging and longevity (Dong et al., 2007; Narayan et al., 2016; Walther et al., 2015). In our studies, we have analyzed WT, *daf-2(lf)*, and *daf-2(lf);daf-16(lf)* worms using five different quantitative proteomics methods: TMT-SPS-MS3, TMT-MS2, LFQ-DIA, LFQ-DDA, and SILAC-spike-in. We also compared the performance of Lumos, Ascend, and Astral mass spectrometers side-by-side using the LFQ-DIA proteomic approach. Together, we have identified 166 protein candidates that showed different levels between WT/*daf-2(lf);daf-16(lf)* and the *daf-2(lf)* mutant, but not between WT and the *daf-2(lf);daf-16(lf)* mutant. Further genetic screening of these candidates identified three proteins whose encoding genes inactivation extended lifespan by more than 10%.

## 2 | METHODS

### 2.1 | *C. elegans* and bacteria strains and maintenance

*C. elegans* strains were maintained at 20 °C on standard nematode growth medium (NGM) plates seeded with *E. coli* OP50. The *C. elegans* strains of Bristol WT (wild type), CB1370 (*daf-2(e1370)*), *daf-2(e1370)*;*daf-16(mgDf50)* and bacteria strain *SLE1: E. coli* [argA, lysA, mcrA, mcrB, IN(rrnD-rrnE)1, lambda-, rcn14::Tn10(DE3 lysogen::lavUV5 promoter -T7 polymerase] from Caenorhabditis Genetics Center (CGC), MN, USA were used in this study.

### 2.2 | Chemicals

Dithiothreitol (DTT), iodoacetamide (IAA), formic acid (FA), and trifluoroacetic acid (TFA), urea, Na_2_HPO_4_:7H_2_O, KH_2_PO_4_, NaCl, NH_4_Cl, magnesium sulfate (MgSO_4_), cholesterol, potassium phosphate monobasic (KH_2_PO_4_), calcium chloride (CaCl_2_), agarose, ammonium hydroxide, ammonium bicarbonate, and other chemicals were purchased from Sigma (St. Louis, MO, USA). Isopropyl-beta-D-thiogalactoside (IPTG), ampicillin, carbenicillin, and tetracycline hydrochloride were purchased from GoldBio (St. Louis, MO, USA). Sequencing Grade Modified Trypsin (Lyophilized) was purchased from Promega (Madison, WI, USA). Lysyl Endoproteinase (Lys-C) was purchased from Wako Chemicals VWR (PA, USA). Optima™ LC/MS Grade water, acetonitrile, methanol, isopropanol were purchased from Fisher Scientific (MA, USA). Sep-Pak tC18 3cc Vac cartridge was purchased from Waters (MA, USA). Biognosys iRT kit was purchased from Bruker BioSpin Corporation (MA, USA). 1 M triethylammonium bicarbonate (TEAB), 50% hydroxylamine, TMT™ 10plex Label Reagent Set, 1.0 M Tris/HCl (pH=8.0, Invitrogen), Pierce™ BCA Protein Assay Kit-Reducing Agent Compatible, Pierce™ Peptide Desalting Spin Columns, Pierce™ Quantitative Colorimetric Peptide Assay, Pierce™ Quantitative Fluorometric Peptide Assay were purchased from ThermoFisher Scientific (MA, USA). Deionized water was prepared by a Milli-Q system (Millipore, Bedford, MA). Heavy-labeled amino acids: Arg10: L-Arginine 13C6, 15N4 hydrochloride (>98% purity), Lys8: L-Lysine U-13C6, 15N2 hydrochloride (>98% purity) were purchased from AthenaES (MD, USA). OmniPur Sodium dodecyl sulfate (SDS) was purchased from EMD Chemicals Inc., Merk (NJ, USA). cOmplete™, EDTA-free Protease Inhibitor Cocktail (SKU 11873580001), PhosSTOP™ (SKU 4906845001) were purchased from Millipore Sigma (MA, USA). 1.0 M Tris/HCl (pH=8.5) was purchased from Rockland (PA, USA).

### 2.3 | Protein extraction and trypsin digestion

Three biological replicates of each condition (WT, *daf-2(lf)*, *daf-2(lf);daf-16(lf)*) were used. The adult Day-1 worms were collected with M9 buffer, washed the worms with M9 buffer generously three times. Worms were lysed in 4% SDT lysis buffer (4% SDS, 100 mM Tris-HCl, 100 mM DTT) with EDTA-free Protease Inhibitor Cocktail to final concentration of 1×, and PhosSTOP™ to final concentration of 1× by homogenized with beats beater. The lysate was ultrasonicated to shear DNA and then centrifuged for 10 min at 13,000 rpm at room temperature, and then the supernatant was collected. A BCA assay was performed to measure protein concentration by using Pierce™ BCA Protein Assay Kit-Reducing Agent Compatible. The filter-aided sample preparation (FASP) method (Wisniewski et al., 2009) was used for protein reduction, alkylation, and LysC (1 mAU/50 µg protein, 2 hours, 37 °C) followed by trypsin (1:50 (w/w), 16 hours, 37 °C) digestion. After getting the peptides, they were desalted by Sep-Pak tC18 Cartridge (200 mg) from Waters. The TMT-labeled peptides were desalted by Pierce™ peptide desalting spin columns. Then the concentration was estimated by using Pierce™ Quantitative Colorimetric/Fluorometric Peptide Assay. The peptides were aliquot to the same concentration of each sample, dried in SpeedVac and stored in -80 °C until use.

### 2.4 | TMT labeling

TMT10 plex labeling was conducted according to manufacturer’s instructions and previous report. After confirming the labeling efficiency, the same amount of each channel was combined, quenched, dried in SpeedVac.

### 2.5 | Stable isotope labeling by amino acid (SILAC) labeling

The stable isotope labeling was performed as described previously with some modifications(Larance et al., 2011). Inoculate SLE1 into M9 minimal medium with Heavy L-arginine and L-lysine. The 3rd culture, grow overnight at 37 °C until OD600=1. Double stranded RNA targeting orn-1 is induced by the addition of IPTG. After egg preparation of worms, the synchronized L1 WT worms were seeded onto 15-cm NGM-noN plates (without nitrogen source) (Table S10) with corresponding SILAC-labeled SLE1 bacteria to adulthood as the first generation, and then egg preparation and synchronization were conducted from the first generation to get the second generation. The labeling efficiency was evaluated for each generation by LC-MS. The labeled and unlabeled protein lysates were equally mixed at the protein level prior to digestion.

### 2.6 | High-pH reverse phase HPLC fractionation

The high-pH reverse phase offline fractionation was conducted for the dried TMT-labeled peptides sample and label-free DDA library-built separately. The sample was resuspended in solvent A (4.5 mM ammonium formate (pH 10) in 2% acetonitrile). The solvent B is composed of 4.5 mM ammonium formate (pH 10) in 90% acetonitrile. The peptides were separated on a Zorbax 300Extend-C18 column (4.6 × 250 mm, 3.5-Micron, Agilent, CA, USA) at flow rate of 1 mL/min, and collected on Agilent collector (G5664B, 1260 Bio FC-AS) with a gradient of 0% B for 0-9 min, then by a linear gradient from 0% B to 6% B in 4 min, followed by a linear increase to 28.5% B in 50 min, 34% B in 5.5 min, 60% in 14 min, and kept there for 8.5 min for label-free DDA library-built sample; 0% B for 0-7 min, then by a linear gradient from 0% B to 16% B in 6 min, followed by a linear increase to 40% B in 60 min, 44% B in 4 min, 60% in 5 min, and kept there for 4 min for the TMT-labeled sample. The 24 fractions were dried in SpeedVac and ready for LC-MS.

### 2.7 | NanoLC−MS/MS analysis on Orbitrap Fusion Lumos

The label-free DIA and DDA, TMT, SILAC-spike-in samples were analyzed on an Orbitrap Fusion Lumos mass spectrometer coupled to an EASY-nLC 1200 system (Thermo Fisher Scientific, USA). iRT was added into the LFQ-DIA method samples based on the manufacture’s introduction. Peptides were separated on a C18 trap column (Acclaim™ PepMap™ 100 C18 HPLC Column, 3 μm, 75 μm × 2 cm, Thermo Fisher Scientific) coupled with an analytical column (Acclaim™ PepMap™ 100 C18 HPLC Column, 2 μm, 75 μm × 15 cm, Thermo Fisher Scientific) using an 130-min, 140-min per fraction, 80-min gradient for label-free, TMT, SILAC-spike-in methods respectively (Table S5). The detailed parameters of MS1, MS2, or MS3 were set as shown in Table S11. For label-free DIA, quadrupole isolation mode was selected with variable isolation windows based on DDA library data and calculated by SWATH Variable Window Calculator (https://sciex.com/support/knowledge-base-articles/how-to-use-the-swath-variable-calculator-excel-sheet_en_us) as shown in Table S2. The flow rate was 300 nL/min, and the solvent A and solvent B were 0.1% FA in H_2_O and 0.1% FA in 80% ACN, respectively.

### 2.8 | NeoLC−MS/MS analysis on Orbitrap Ascend

Ascend-DIA samples were analyzed on an Orbitrap Ascend mass spectrometer coupled to an Vanquish™ Neo UHPLC System (Thermo Fisher Scientific, USA). iRT was added into the samples based on the manufacture’s introduction. Peptides were separated on a C18 trap column (Acclaim™ PepMap™ 100 C18 HPLC Column, 3 μm, 75 μm × 2 cm, Thermo Fisher Scientific) coupled with an analytical column (Easy-Spray™ PepMap™ Neo 2 μm C18 75 μm × 150 mm, Thermo Fisher Scientific and Column: Aurora Ultimate XT 25 × 75 C18 UHPLC column, 25 cm × 75 μm, 1.7 μm) using an 130 min (Table S5). MS1 scans were collected at 120K resolution, with an AGC target of 1 × 10^6^ and maximum injection time of 251 ms. Quadrupole isolation mode was selected with an isolation window of 14 m/z. Fragmentation was performed using an HCD activation energy of 28%. AGC target in MS2 was set to 5 × 10^5^, maximum injection time of 59 ms and the analyzer to Orbitrap with a resolution at 30K. The precursor mass range was set as 400-1040 m/z.

### 2.9 | NeoLC−MS/MS analysis on Orbitrap Astral

Astral-DIA samples were analyzed on an Orbitrap Astral mass spectrometer coupled to an Vanquish™ Neo UHPLC System (Thermo Fisher Scientific, USA). iRT was added into the samples based on the manufacture’s introduction. Peptides were separated on a C18 column (PepMap™ Neo Trap Cartridge with PepMap™ Neo UHPLC Column, 5mm, Thermo Fisher Scientific) and an analytical column (EASY-Spray™ HPLC Columns, 15 cm × 150 μm, 2 μm, Thermo Fisher Scientific) using a 24.4 min (Table S5). MS1 scans were collected at 240K resolution, with an AGC target of 5 × 10^6^ and time of 5 ms. Quadrupole isolation mode was selected with an isolation window of 2 m/z. Fragmentation was performed using an HCD activation energy of 25%. AGC target in MS2 was set to 5 × 10^4^ and time of 5 ms and the analyzer to Astral. The precursor mass range was set as 380-980 m/z.

### 2.10 | Protein identification, quantification, and statistical analysis

The raw files of LFQ-DDA, TMT-MS2/MS3, and SILAC-spike-in were analyzed with MSFragger (version 4.1) in FragPipe (version 22.0)(Djomehri et al., 2020; Kong et al., 2017; Teo et al., 2021) and searched against UniprotKB *C. elegans* reference database, downloaded on December 18, 2023 (27,454 entries). General settings include dynamic modifications: oxidation (+15.995 Da) on methionine, acetyl (protein N-term). The fixed modifications: carbamidomethyl (+57.021 Da) on cysteine. The allowing missed cleavage was set to 2. The precursor and fragment mass tolerance were set to 20 ppm.

For TMT-labeled data, the workflow of TMT10 and TMT10-MS3 were used for TMT-MS2 and TMT-MS3 respectively. The TMT (+229.163 Da) on peptide N-terminus and lysine was set as a static modification. The other parameters were set as default.

For LFQ-DDA data, the workflow of LFQ-MBR (label-free quantification-match between runs) was used. The label minimum ratio count was set as 2 for quantification using MaxLFQ. The other parameters were set as default.

For SILAC-spike-in data, the workflow of SILAC3 was selected with slight modification. The isotope labeling of lysine (+8.012), arginine (+10.008), and proline (+6.014) were enabled as variable modifications. The other parameters were set as default.

The raw files of TMT labeling efficiency and peptides fractionation of LFQ-DDA for library-based DIA method optimization on Lumos were analyzed with Proteome Discoverer 2.4 and MaxQuant (version 2.1) using MaxDIA(Sinitcyn et al., 2021) respectively.

The raw files of LFQ-DIA were analyzed with DIA-NN (version 1.8.1) and searched against UniprotKB *C. elegans* reference database, downloaded on December 18, 2023 (27,454 entries) and the Biognosys iRT peptides fasta(Bruderer et al., 2016). General settings include dynamic modifications: oxidation (+15.995 Da) on methionine. The fixed modifications: carbamidomethyl (+57.021 Da) on cysteine. The protease trypsin was selected and the allowing missed cleavage was set to 2. The minimum and maximum peptide length were set to 6 and 40 respectively. The precursor range (m/z) was set at 300-2000, and the precursor charge was set at 2-6. The precursor and fragment mass tolerance were set to 10 ppm. Match between runs (MBR) was enabled. Deep learning was used to generate a new in silico spectral library from peptides provided. Library precursor was reannotated using the FASTA database. Library-free search was enabled. The precursor FDR was set to 1%. The scan window radius was set to 6. Library generation was set to IDs, RT and IM profiling. The neural network classifier was set to double-pass mode. The quantification mode was set to robust LC (high precision). All other settings were left default.

All output quantification data, corresponding to protein-encoding genes names, from MSFragger and DIA-NN were log_2_-transformed and filtered to retain at least two valid values in at least one group. Within-condition reproducibility was assessed using protein-group coefficients of variation (CVs) calculated from linear abundance values across biological replicates within each condition. For each workflow and condition, CV was calculated as the standard deviation divided by the mean abundance across biological replicates and expressed as a percentage. For each workflow, we summarized the CV distribution by median, interquartile range (IQR), and the proportion of protein groups with CV below 10%, 20%, and 30%. To control for differences in workflow-dependent protein coverage, we repeated the analysis using condition-specific common protein group sets. Missing values were imputed using a normal distribution (width: 0.3 standard deviation; down shift: 1.8 standard deviation) and principal component analysis (PCA) was subsequently performed using Perseus(Tyanova et al., 2016). Then, DEqMS (Differential Expression analysis of quantitative Mass Spectrometry) (Zhu et al., 2020), a statistical method designed to identify differentially expressed proteins in quantitative proteomics, was performed to identify the significantly differentially expressed proteins. The peptide counts of LFQ-DIA method were combined from the output from DIA-NN. DEqMS was used for significance analysis at a significance level of 0.05 with adjusted DEqMS p-values (sca.adj.pval). At least two-fold changed proteins with significance were considered as differentially expressed proteins. The UpSet plot was generated using R.(Conway et al., 2017) The datasets were merged and filtered using Python. The volcano plot, Pearson’s correlation and survival curve were generated using GraphPad Prism 10. Figures and graphs were constructed using BioRender and Illustrator.

### 2.11 | LFQ-DIA-Astral-referenced recovery analysis

We performed an Astral-centered recovery analysis to evaluate how well more broadly accessible proteomics workflows reproduced LFQ-DIA-Astral-derived candidate lists, rankings, and direction-of-change patterns. LFQ-DIA-Astral DE candidates were used as the reference set for each comparison and regulation direction. Candidate proteins were ranked by LFQ-DIA-Astral log2 fold change: up-regulated candidates were ranked from largest positive log2FC to smaller positive log2FC, whereas down-regulated candidates were ranked from most negative log2FC to less negative log2FC. For each non-Astral workflow, the corresponding all-protein table was used, so that recovery represented whether an Astral candidate was quantified and testable in that workflow. For each Astral candidate and each workflow, three recovery metrics were calculated separately. Detection recovery: the Astral candidate was present/quantified in the workflow’s all-protein table. Direction concordance: the Astral candidate was detected and had the same log2FC sign in the workflow as in LFQ-DIA-Astral. Strict statistical support: the Astral candidate was detected, had the same log2FC sign as Astral DIA, and had sca.adj.pval ≤ 0.05 in the workflow. We then evaluated all single workflows and all possible workflow combinations among LFQ-DIA-Ascend, LFQ-DIA-Lumos, LFQ-DDA, TMT-MS2, and TMT-SPS-MS3 (Peng et al., 2024;

Sissala et al., 2025). For a workflow combination, a candidate was counted as recovered if at least one workflow in the combination satisfied the metric being evaluated. Thus, combination-level detection recovery required detection by any workflow in the combination, combination-level direction concordance required same-direction log2FC in any workflow in the combination, and combination-level strict statistical support required same-direction log2FC and sca.adj.pval ≤ 0.05 in any workflow in the combination. For cumulative recovery plots, Astral candidates were traversed in ranked order. At each rank threshold *k*, we calculated the percentage of the top *k* Astral candidates that were recovered by each workflow or workflow combination under the relevant metric. Separate cumulative plots were generated for detection recovery, direction concordance, and strict statistical support. These plots therefore show how rapidly each workflow or workflow combination recovers the highest-ranked LFQ-DIA-Astral DE candidates as progressively lower-ranked Astral candidates are included.

### 2.12 | Functional analysis of differentially expressed proteins

The Wormbase ID of each gene was retrieved using WormMine (https://intermine.wormbase.org/tools/wormmine/), part of the WormBase project. The WormCat was used for functional analysis of differentially expressed proteins under classification in physiological (Category 1), molecular (Category 2), or subcellular localization (Category 3) categories.(Holdorf et al., 2020)

### 2.13 | RNA interference (RNAi) experiments

RNAi libraries created by the lab of Dr. Marc Vidal and Dr. Julie Ahringer were used in this study.(Kamath & Ahringer, 2003; Rual et al., 2004) The RNAi clones of cat-4, K10D3.4, wrt-1, and taf-2 were acquired from the Vidal library while agr-1, F44E2.4 and clec-10 RNAi clones were acquired from the Ahringer library. All RNAi clones were verified by Sanger sequencing. All RNAi colonies were selected in both 100 μg/mL ampicillin and 15 μg/mL tetracycline resistance. All RNAi bacteria were cultured for 14 hours in LB with 60 μg/mL carbenicillin and then seeded onto RNAi agar plates that contain 1 mM IPTG and 25 μg/mL carbenicillin. The plates were then left at room temperature overnight for induction of dsRNA expression.

### 2.14 | Lifespan assays

The WT worms were synchronized by bleach-based egg preparation and subsequent starvation in M9 buffer overnight. Synchronized L1 larvae were plated onto 6-cm freshly made RNAi plates seeded with the bacteria that carry specific RNAi clones. The animals were synchronized again by manual picking at mid L4 stage with 30-40 worms per plate and 60-120 worms per condition. and marked as day 0. Adult worms were observed and transferred to new freshly made RNAi plates every day or every other day. Animals were categorized as alive, dead (cessation of movement in response to platinum wire gentle probing), or censored. The statistical analyses were performed using IBM SPSS Statistics 29.0 with Kaplan–Meier survival function and the log-rank test. GraphPad Prism 9 was used to graph the results. Three independent biological replicates. For all figure legends, asterisks indicate statistical significance as follows: n.s. = not significant p>0.05; * p<0.05; ** p<0.01; *** p<0.001.

## 3 | RESULTS

### 3.1 | Comparative proteomics between long- and normal-lived animals across methods

We have applied five different quantitative MS proteomic approaches to compare the proteomes among three biological groups of *C. elegans*: wild type (WT), the long-lived *daf-2(lf)* mutant, and the normal-lived *daf-2(lf);daf-16(lf)* double mutant. These include tandem-mass-tags (TMT) isobaric labeling-based proteomics at both MS2 and MS3 level, label-free quantification (LFQ) with data-dependent-acquisition (DDA) or data-independent acquisition (DIA)-based proteomics, and stable isotope labeling with amino acid (SILAC) spike-in-assisted proteomics. Using LFQ-DIA, we also compared three versions of Orbitrap mass spectrometers: Fusion Lumos, Ascend, and Astral (Figure 1).

**Figure 1.**
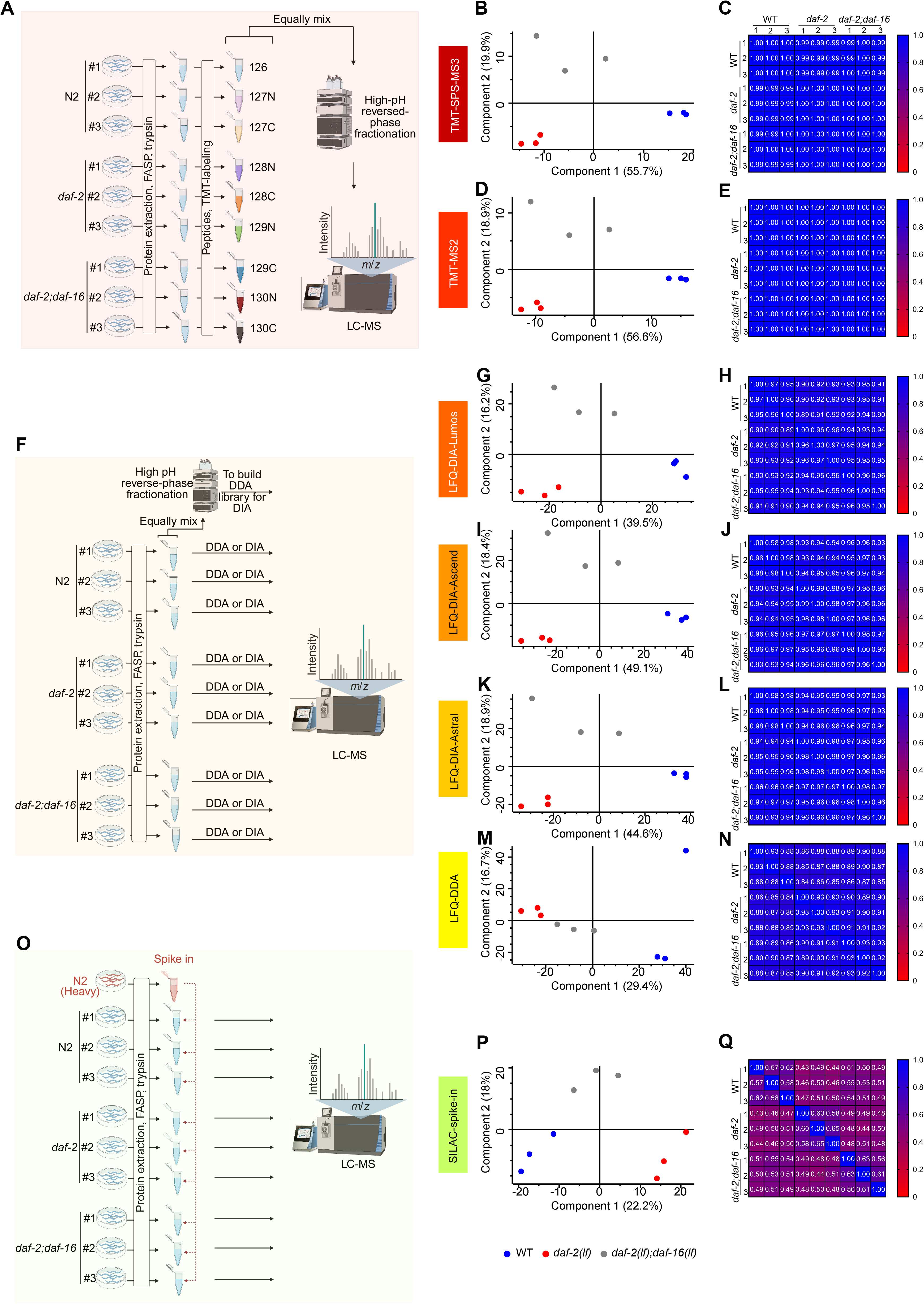
Summary of quantitative proteomics using different methods. (A) Workflow of tandem mass tag (TMT)-based proteomics. (B) Principal component analysis (PCA) of TMT-SPS-MS3 shows that three genotypic groups (WT, *daf-2(lf)*, *daf-2(lf);daf-16(lf)*) form distinct clusters with Component 1 accounting for 55.7% and Component 2 for 19.9% of total variance. (C) Heatmap of Pearson’s correlation coefficients of TMT-SPS-MS3 among three biological replicates from WT, *daf-2(lf)*, and *daf-2(lf);daf-16(lf)* shows coefficient r > 0.98. (D) PCA of TMT-MS2 shows that three genotypic groups (WT, *daf-2(lf)*, *daf-2(lf);daf-16(lf)*) form distinct clusters with Component 1 accounting for 56.6% and Component 2 for 18.9% of total variance. (E) Heatmap of Pearson’s correlation coefficients of TMT-MS2 among three replicates from WT, *daf-2(lf)*, and *daf-2(lf);daf-16(lf)* shows coefficient r > 0.98. (F) Workflow of label-free quantification (LFQ)-based proteomics. (G) PCA of LFQ-DIA-Lumos shows that three genotypic groups (WT, *daf-2(lf)*, *daf-2(lf);daf-16(lf)*) form distinct clusters with Component 1 accounting for 39.5% and Component 2 for 16.2% of total variance. (H) Heatmap of Pearson’s correlation coefficients of LFQ-DIA-Lumos among three replicates from WT, *daf-2(lf)*, and *daf-2(lf);daf-16(lf)* shows coefficient r > 0.88. (I) PCA of LFQ-DIA-Ascend shows that three genotypic groups (WT, *daf-2(lf)*, *daf-2(lf);daf-16(lf)*) form distinct clusters with Component 1 accounting for 49.1% and Component 2 for 18.4% of total variance. (J) Heatmap of Pearson’s correlation coefficients of LFQ-DIA-Ascend among three replicates from WT, *daf-2(lf)*, and *daf-2(lf);daf-16(lf)* shows coefficient r > 0.92. (K) PCA of LFQ-DIA-Astral shows that three genotypic groups (WT, *daf-2(lf)*, *daf-2(lf);daf-16(lf)*) form distinct clusters with Component 1 accounting for 44.6% and Component 2 for 18.9% of total variance. (L) Heatmap of Pearson’s correlation coefficients of LFQ-DIA-Astral among three replicates from WT, *daf-2(lf)*, and *daf-2(lf);daf-16(lf)* shows coefficient r > 0.92. (M) PCA of LFQ-DDA shows that three groups (WT, *daf-2(lf)*, *daf-2(lf);daf-16(lf)*) form distinct clusters with Component 1 accounting for 29.4% and Component 2 for 16.7% of total variance. (N) Heatmap of Pearson’s correlation coefficients of LFQ-DDA among three replicates from WT, *daf-2(lf)*, and *daf-2(lf);daf-16(lf)* shows coefficient r > 0.87. (O) Workflow of stable isotope labeling by amino acids (SILAC)-spike-in-based proteomics. (P) PCA of SILAC-spike-in shows that three genotypic groups (WT, *daf-2(lf)*, *daf-2(lf);daf-16(lf)*) form distinct clusters with Component 1 accounting for 22.2% and Component 2 for 18.0% of total variance. (Q) Heatmap of Pearson’s correlation coefficients of SILAC-spike-in among three replicates from WT, *daf-2(lf)*, and *daf-2(lf);daf-16(lf)* shows coefficient 0.55 < r < 0.65. Pearson’s correlation: color intensity represents correlation strength from +1 (strong positive) to 0 (no correlation). Cells are annotated with correlation r values.

In the TMT method, peptides derived from each biological sample were chemically labeled with distinct tandem mass tags after digestion, then pooled and fractionated using high-pH reverse-phase chromatography (Figure 1A). The labeling efficiency was evaluated before fractionation and exceeded 98% for both lysine and N-terminus (Table S1). Using the Orbitrap Fusion Lumos, both MS2 and MS3 quantifications were conducted to enhance quantification accuracy and reduce ratio distortion. PCA analysis of candidates identified by the TMT method showed that three replicates from each biological group clustered together and were clearly separated from other groups, indicating that TMT effectively captured underlying biological variability despite technical multiplexing. The first two principal components accounted for 75.6% and 75.5% of the total variance in SPS-MS3 and MS2 respectively, with PC1 separating WT and *daf-2(lf)*, and PC2 distinguishing between *daf-2(lf);daf-16(lf)* and WT or *daf-2(lf)* (Figure 1B and 1D). This clustering pattern aligns with lifespan differences among these three biological groups, highlighting the robustness of the TMT method in distinguishing biological states. Furthermore, Pearson’s correlation analysis demonstrated very strong positive correlations (coefficient r > 0.99) among replicates, underscoring high reproducibility (Figure 1C and 1E).

For the LFQ method, both DIA and DDA strategies were employed. DIA with library-free approaches to enhance peptide identification and quantification across samples (Figure 1F). Specifically, we first utilized an LFQ-DDA for fractioned peptides to build the library to optimize acquisition parameters for DIA approach on the Orbitrap Lumos instrument (Table S2), followed by a library-free strategy for final quantification. For samples analyzed on the Orbitrap Ascend and Orbitrap Astral instruments, the library-free DIA approach was applied using the optimized parameters (Table S3). We also evaluated the performance of different columns (Thermo Fisher Scientific Easy-Spray™ PepMap™ (15 cm) and Ion Opticks (IO) Aurora Ultimate (25 cm) C18 columns) in DIA analysis using Orbitrap Ascend. We found that although IO column enabled identification of more proteins within the same gradient, the average data points of each peak were less than 6, even when the isolation window was increased from 14 to 18 Th (Figure S1). Although a longer gradient may improve performance on the IO column, we selected the 15 cm column as the primary choice in our studies, as it was also consistent with all the other methods.

PCA analysis of the candidates identified using the LFQ-DIA method were able to separate the three distinct biological groups, albeit with slightly higher technical variance compared to TMT. The first two principal components accounted for 55.7%, 67.5%, and 63.5% of the total variance when using Orbitrap Fusion Lumos (Figure 1G), Orbitrap Ascend (Figure 1I), and Orbitrap Astral (Figure 1K) instruments, respectively. PC1 primarily separated WT and *daf-2(lf)*, while PC2 distinguished *daf-2(lf);daf-16(lf)* from both WT and *daf-2(lf)*. Pearson’s correlation analysis demonstrated very strong positive correlations (coefficients r > 0.95, r > 0.97, r > 0.97) between replicates, underscoring high reproducibility (Figure 1H, 1J, and 1L). For the candidates identified using LFQ-DDA on the Orbitrap Fusion Lumos, PCA analysis distinguished *daf-2(lf)* from WT and *daf-2(lf);daf-16(lf)* (Figure 1M), while Pearson’s correlation analysis showed strong positive correlations among replicates (coefficient r > 0.88) (Figure 1N). Although this method yielded lower overall performance compared to LFQ-DIA, its reproducibility remains acceptable.

In the SILAC-spike-in proteomic workflow (Figure 1O), SLE1 bacteria were first labeled by culturing them with stable isotope-labeled lysine and arginine and then fed to *C. elegans* to achieve isotopic labeling. We confirmed that the labeling efficiency in WT *C. elegans* exceeded 97% after two-generations of growth on isotope-labeled bacteria (Table S4). For spike-in quantification, labeled and unlabeled protein lysates were mixed at the protein level prior to digestion, and the resulting peptide mixtures were analyzed using the Orbitrap Fusion Lumos. PCA analysis of the candidates identified using the SILAC-spike-in workflow showed clear clustering of samples according to their biological groups (Figure 1P). The first two principal components accounted for 61.3% of the total variance, with PC1 separating WT and *daf-2(lf)* and PC2 distinguishing *daf-2(lf);daf-16(lf)* from WT or *daf-2(lf)* (Figure 1P). However, Pearson’s correlation analysis showed moderate correlations (coefficient r > 0.56) between replicates, indicating less reproducibility compared to other methods (Figure 1Q). In addition, across all quantified protein groups, median within-condition CVs were below 30% for every workflow and condition, indicating strong within-condition quantitative precision across workflows, while the common-set analysis confirms that the observed workflow differences are not solely driven by method-dependent protein coverage. Quantitative precision was highest for TMT-MS2 and TMT-SPS-MS3, followed by LFQ-DIA-Astral, LFQ-DIA-Ascend, and LFQ-DIA-Lumos, with LFQ-DDA showing the greatest replicate variability (Figure S2).

Together, these systematic side-by-side comparisons suggest that TMT and LFQ-DIA offer higher precision and reproducibility in capturing biological differences, along with higher overall proteome coverage.

### 3.2 | Identification comparison of longevity-associated proteins across proteomic analyses

Next, we comprehensively compared the numbers of identified peptides, identified proteins, and quantifiable proteins, non-missing value rates, as well as differentially expressed (DE) proteins between groups across the different proteomic methods and the different MS instruments (Figure 2). TMT (with peptides fractionation), LFQ-DDA, LFQ-DIA and SILAC-spike-in analyses on the Orbitrap Fusion Lumos were first assessed. In terms of workflow-level identification depth, fractionated TMT-MS2 yielded the largest number of identifications, with approximately 70,000 peptides and 8,000 protein groups. Fractionated TMT-SPS-MS3 identified substantially fewer peptides (∼40,000, ∼43% decrease) and moderately fewer protein groups (∼6,000, ∼25% decrease) compared with TMT-MS2, consistent with additional MS3 acquisition step reducing MS2 sampling depth and sensitivity. Both TMT workflows identified more protein groups than the LFQ-DDA (∼40,000 peptides and ∼4,000 protein groups) and LFQ-DIA (∼40,000 peptides and ∼5,000 protein groups) workflows under the condition tested. However, this comparison reflects the combined effects of acquisition strategy, multiplexing, and peptide fractionation rather than acquisition mode alone. SILAC-spike-in showed the lowest identification depth in this dataset, with ∼20,000 peptides and 3,000 protein groups (Figure 2A and 2B). This difference became even more profound when assessing the numbers of quantifiable proteins among these methods: ∼6,000 in TMT-SPS-MS3, ∼8,000 in TMT-MS2, ∼5000 in LFQ-DIA, <4,000 in LFQ-DDA, and ∼2,000 in SILAC-spike-in (Figure 2C). Furthermore, the TMT methods exhibited zero missing values, while the non-missing value rates were lower using LFQ-DIA (90%), LFQ-DDA (80%) and SILAC-spike-in (80%) (Figure 2D). It is also worth noting that reagent costs for TMT and isotopically labeled amino acids are substantially higher than for non-labeling methods, and peptide fractionation associated with TMT significantly increases instrument runtime.

**Figure 2.**
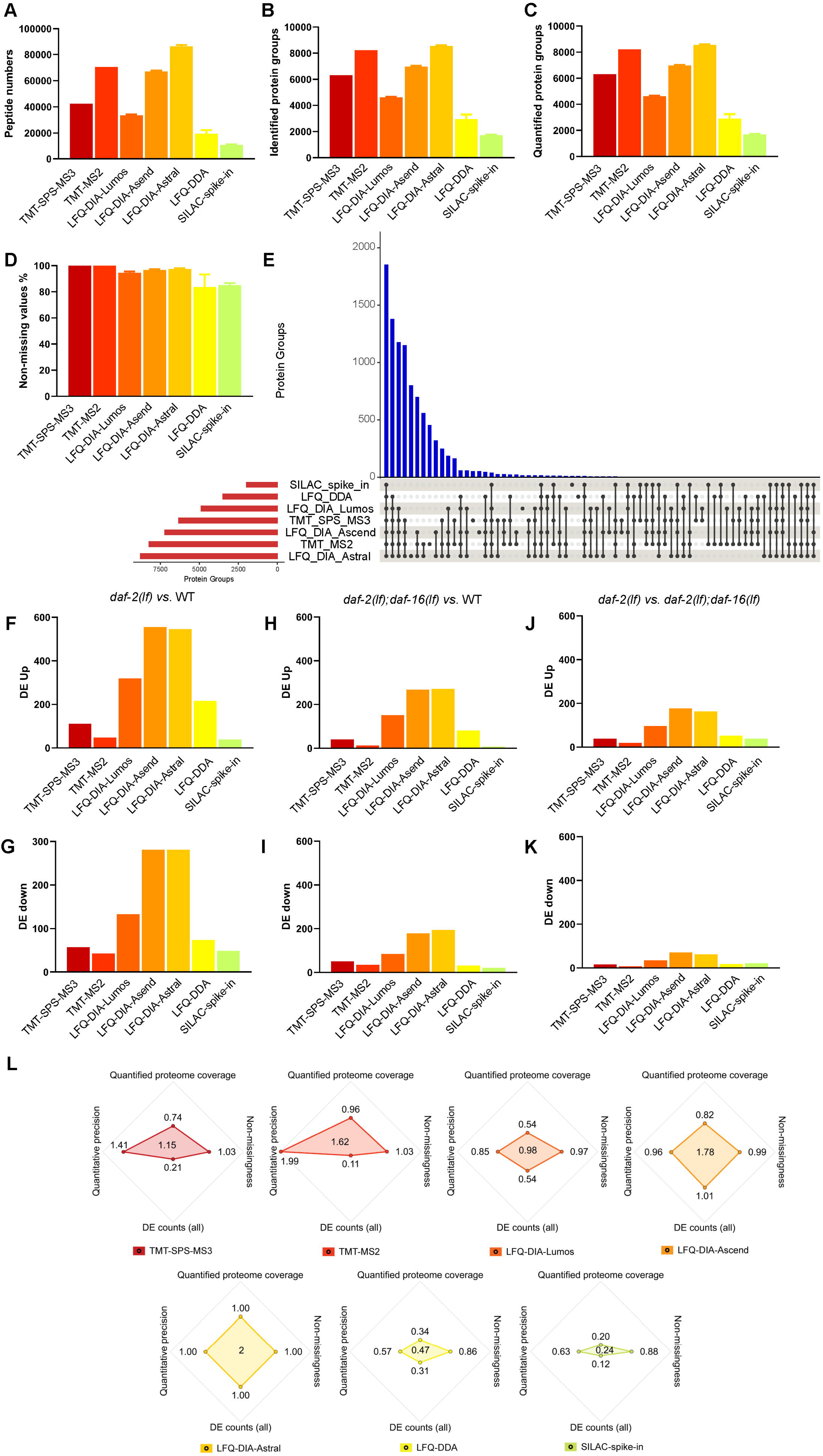
Comparison and integration of different methods in protein identification and quantification. (A) Comparison of identified peptide numbers among different methods. (B) Comparison of identified protein groups among different methods. (C) Comparison of quantitative protein groups among different methods. (D) Comparison of percentage of non-missing values among different methods. (E) UpSet plot shows overlap of quantitative protein groups among different methods. (F) Comparison of differentially expressed protein groups showing up-regulation (DE up) in *daf-2(lf)* vs. WT among different methods. (G) Comparison of differentially expressed protein groups showing down-regulation (DE down) in *daf-2(lf)* vs. WT among different methods. (H) Comparison of DE up protein groups in *daf-2(lf);daf-16(lf)* vs. WT among different methods. (I) Comparison of DE down protein groups in *daf-2(lf);daf-16(lf)* vs. WT among different methods. (J) Comparison of DE up protein groups in *daf-2(lf)* vs. *daf-2(lf);daf-16(lf)* among different methods. (K) Comparison of DE down protein groups in *daf-2(lf)* vs. *daf-2(lf);daf-16(lf)* among different methods. (L) Radar plots summarize proteomics platform performance, separate radar plots show the relative performance profile of each workflow across four metrics: quantified proteome coverage, non-missingness, total differentially expressed protein counts, and average median CV-based quantitative precision. Values were plotted on a shared radial scale from 0 to 2 relative to the reference of LFQ-DIA-Astral for each metric, enabling direct visual comparison across workflows. Larger radial values indicate stronger performance. For the median CV metric, values are represented so that higher values correspond to lower median CV and therefore better quantitative precision. (Bars in Figure 2A-D represent mean values across individual samples and error bars indicate standard deviation (SD). DE: at least two-fold change with adjusted p value ≤ 0.05).

We further compared the performance of three different MS instruments based on LFQ-DIA analyses. We found that the Orbitrap Astral outperformed both Ascend and Lumos in the number of identified peptides (Lumos: 40472, Ascend: 74884, Astral: 93564), identified protein groups (Lumos: 4937, Ascend: 7236, Astral: 8819), and quantified protein groups (Lumos: 4886, Ascend: 7220, Astral: 8783) (Figure 2A-2C). The non-missing value rates were comparable between Ascend (96.73%) and Astral (97.37%) and higher than Lumos (92.49%) (Figure 2D). Notably, Ascend employed a 130-min gradient, whereas Astral used a 24.4-min gradient (Table S5). As such, for high throughput applications, Astral is clearly the most favorable choice. However, when runtime is not a limiting factor, Ascend can approach Astral’s performance with careful optimization and extended acquisition time.

Overall, a substantial number of proteins were consistently identified (3730 protein groups) and quantified (1853 protein groups) across all the methods (Figure S3, Figure 2E). Without including low-performance SILAC-spike-in, the other six methods contributed to the second large overlap in quantification (3232 protein groups) (Figure 2E, total 5085). We further performed differential analysis using DEqMS (Zhu et al., 2020) on each pairwise comparison (*daf-2(lf)* vs. WT, *daf-2(lf) vs*. *daf-2(lf);daf-16(lf)*, *daf-2(lf);daf-16(lf) vs*. WT) to identify significantly altered proteins (adjusted p-value (sca.adj.pval) ≤ 0.05, absolute log_2_ fold change ≥1) (Figure S4). We found that LFQ-DIA using Ascend and Astral identified a relatively higher number of differentially expressed protein candidates between each pairwise comparison, likely due to the higher overall proteome coverage achieved by the LFQ-DIA method on newer-generation of MS instruments (Figure 2F-2K). To our surprise, although TMT-MS2 exhibited high overall proteome coverage, it identified relatively fewer differentially expressed proteins compared to the LFQ-DIA method (Figure 2F-2K). Consistent with prior reports, this difference may be attributed to ratio compression, a well-known common issue in the TMT-MS2 method (Ting et al., 2011). Supporting this, we found that the TMT-SPS-MS3 method—designed to reduce the ratio compression issue—identified more differentially expressed protein than TMT-MS2, despite its lower overall proteome coverage (Figure 2F-2K). We further evaluated the cross-method performance using integrated metrics: quantitative proteome coverage, non-missingness, all differentially expressed protein counts (DE counts (all)), and average median CV-based quantitative precision. For quantitative proteome coverage, non-missingness, and DE counts (all) values from each method were normalized to the corresponding LFQ-DIA-Astral value. For median CV (average), where lower values indicate better quantitative reproducibility, the normalized score was calculated as the ratio of LFQ-DIA-Astral average median CV to the average median CV of each comparison method. Thus, LFQ-DIA-Astral was assigned a normalized value of 1 for each metric, corresponding to a radar-plot polygon area of 2. Using this descriptive composite score, the overall workflow ranking was LFQ-DIA-Astral, LFQ-DIA-Ascend, TMT-MS2, TMT-SPS-MS3, LFQ-DIA-Lumos, LFQ-DDA, and SILAC-spike-in, with radar-plot area scores of 2, 1.78, 1.62, 1.15, 0.98, 0.47, and 0.24, respectively (Figure 2L). Using the same set of biological samples, our analyses directly compare the performance of various comparative proteomic workflows across multiple aspects, including biological reproducibility, overall proteome coverage, identification of differentially expressed protein candidates, cost, and runtime (Table 1). This information provides a reference for selecting proteomic pipelines that best align with specific experimental goals and practical constraints.

**Table 1.** Comparative summary of key features across methods.

| Method | Signal for quantification | Quantitative accuracy | Proteome coverage | Reagent cost | Runtime | Additional note |
| --- | --- | --- | --- | --- | --- | --- |
| TMT-SPS-MS3 | MS3 | + + + + | + + | + + + + | + + + + |  |
| TMT-MS2 | MS2 | + + | + + + + | + + + + | + + + + |  |
| LFQ-DIA-Lumos | MS2 | + + | + + | + | + + + | Require more optimization |
| LFQ-DIA-Ascend | MS2 | + + + | + + + | + | + + + | Require more optimization |
| LFQ-DIA-Astral | MS2 | + + + + | + + + + | + | + + |  |
| LFQ-DDA | MS1 | + + | + | + | + + + |  |
| SILAC-spike-in | MS1 | + | + | + + + + | + + + | Laborious sample preparation |

For further biological analysis based on differentially expressed proteins between conditions, their identification consistency among biological replicates is paramount (Cairns, 2011). The lower Pearson’s correlation coefficient of the SILAC-spike-in dataset indicates higher variability, which could introduce false positive or exclude false negative candidates. Several factors may contribute to this reduced reproducibility, including the sensitivity of ratio-based quantification to noise at low protein abundance, possible partial mismatch between the heavy-labeled reference and the compared genotypes, and other possible factors such as imputation of missing values and stochastic peptide sampling. Given this uncertainty, we conservatively did not include this dataset for the integrative differential expression analysis and focused on the other six datasets by crossing validating their candidates for each other. Specifically, we first integrated the differentially expressed proteins between the long-lived *daf-2(lf)* mutant and WT from each method (the absolute log_2_FC ≥ 1, adjusted p value ≤ 0.05.), which includes 959 up-regulated and 568 down-regulated candidates in the long-lived state (Table S6 (Excel sheet 1)). In comparison, similar analysis of the long-lived *daf-2(lf)* mutant versus the normal-lived *daf-2(lf);daf-16(lf)* double mutant uncovered 271 and 121 upregulated and downregulated proteins, respectively (Table S6 (Excel sheet 2)). Second, to further enhance confidence, we mapped these integrated candidates back to each dataset to obtain the fold change and significance of each candidate across methods. Theoretically, there exists a “true” ratio of the molecular level for each protein candidate between conditions. However, this ratio in real-world proteomic measurements is complicated by numerous technical factors, including ionization efficiency, labeling chemistry, and instrument acquisition strategy, which were handled differently in each method.(Barkovits et al., 2020; Fernandez-Costa et al., 2020; Liu et al., 2007; Wang et al., 2024; Zhang et al., 2013) As a result, no single method would likely capture the “true” ratio, but each one would provide a proximation (Bantscheff et al., 2012; Bantscheff et al., 2007). Consistently, when integrating results from six different methods, we noticed that most of the candidates showed consistent trend change, which supports their reproductivity; however, their exact ratios vary across methods. Therefore, we assigned three scores to each candidate in each method, not detected (N), significant (1), nonsignificant (0), and two scores to each candidate across methods, consistent trend of change (C), inconsistent trend of change (I) (Table S7). The candidates that reached (C, 1 or N) are our first-class confident candidates, specifically, first-class confident candidates require a consistent trend across methods and significant support in at least two methods.

Together, we identified 442 and 216 candidates that are differentially up- and down-regulated, respectively, in the *daf-2(lf)* mutant compared to WT (Table S6 (Excel sheet 3)). Similarly, we identified 112 and 36 candidates that are differentially up- and down-regulated, respectively, in the *daf-2(lf)* mutant compared to the normal-lived *daf-2(lf);daf-16(lf)* double mutant (Table S6 (Excel sheet 4)).

After comparing candidate lists across workflows, we observed incomplete overlaps among methods, indicating that the candidate space was method dependent. Because Astral-DIA yielded the broadest overall peptide and protein-group coverage, we used Astral-DIA as a high-coverage reference to evaluate how well alternative workflows recovered Astral-supported candidates and their direction-of-change patterns.

For each pairwise genotype comparison, LFQ-DIA-Astral upregulated and downregulated differentially expressed (DE) candidates were analyzed separately against each alternative method or method combination. We calculated detection recovery, direction concordance and strict statistical support among shared candidates. Detection recovery measured whether LFQ-DIA-Astral DE candidates were quantified by the comparison workflow. Direction concordance measured the percentage of shared candidates with the same log2FC direction. Strict statistical support measured the subset of LFQ-DIA-Astral DE candidates that were also significant and regulated in the same direction by the comparison workflow (Figure 3, Figure S5-7).

**Figure 3.**
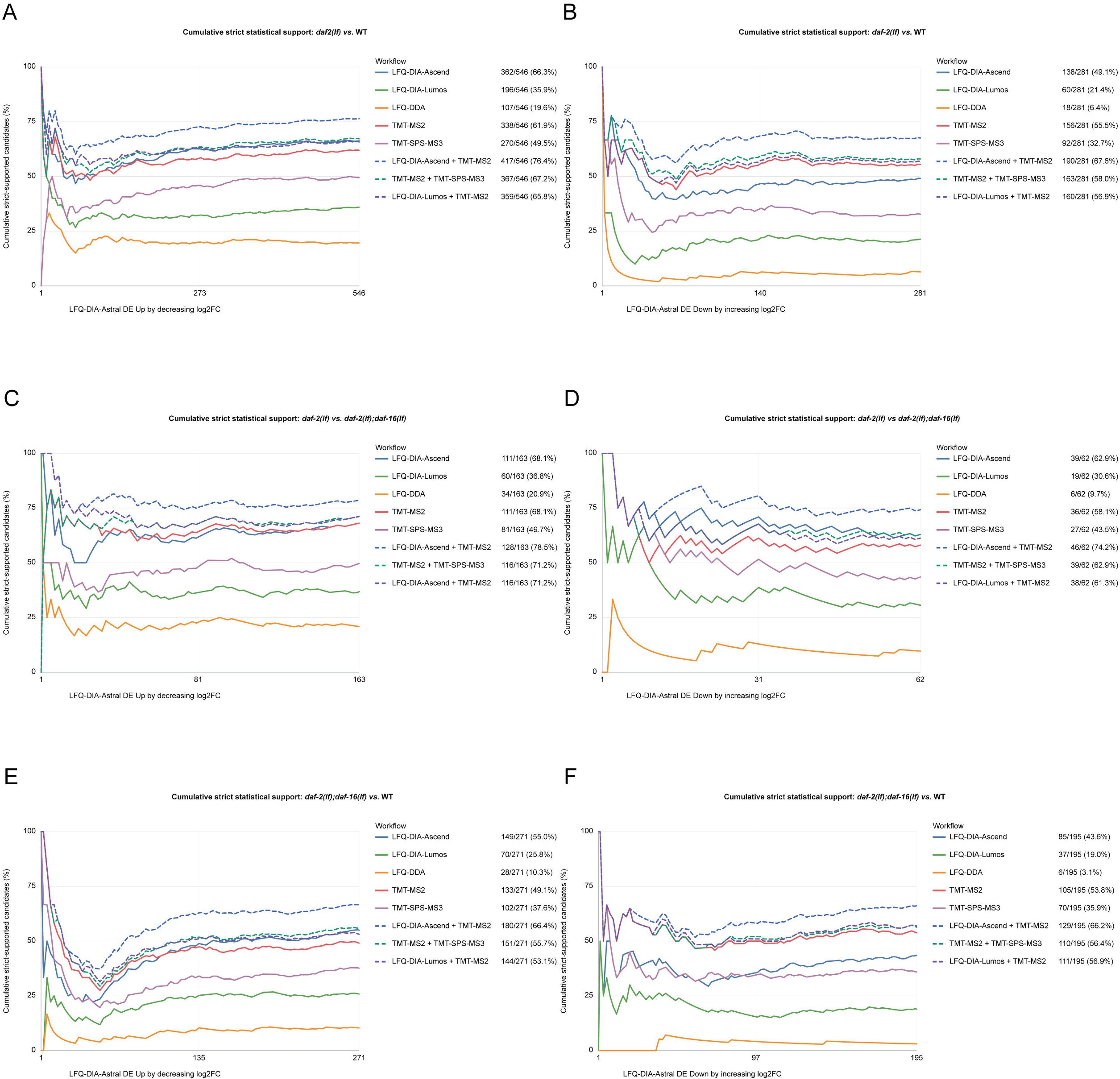
Detection recovery of LFQ-DIA-Astral DE candidates by alternative workflows. Cumulative recovery curves show the proportion of Astral DIA-ranked differentially expressed protein candidates that were quantified or detected by each comparison workflow or workflow combination. Astral candidates were ranked by Astral DIA effect size, with up-regulated candidates ordered by decreasing log2FC and down-regulated candidates ordered by increasing log2FC. (A) Up-regulated DE candidates between *daf-2(lf)* and WT. (B) Down-regulated DE candidates between *daf-2(lf)* and WT. (C) Up-regulated DE candidates between *daf-2(lf)* and *daf-2(lf);daf-16(lf)*. (D) Down-regulated DE candidates between *daf-2(lf)* and *daf-2(lf);daf-16(lf)*. (E) Up-regulated DE candidates between *daf-2(lf);daf-16(lf)* and WT. (F) Down-regulated DE candidates between *daf-2(lf);daf-16(lf)* and WT.

No single non-Astral method or method combination recovered 100% of LFQ-DIA-Astral DE candidates, indicating that the workflows are not fully interchangeable. The combination of all other five workflows provides the maximum recovery (Figure S5). However, the combination of LFQ-DIA-Ascend and TMT-MS2 can reach similar recovery rates at ∼86.3% of detection recovery, ∼82.7% of direction concordance, ∼71.6% of strict statistical support across the three pairwise genotype comparisons. Practically consider the availability of instrumentation, single workflow LFQ-DIA-Ascend has recovery rates at ∼75.5% of detection recovery, ∼73.1% of direction concordance, ∼57.5% of strict statistical support across the three pairwise genotype comparisons (Figure 3). Among Lumos-accessible workflows, the combination of Lumos-DIA and Lumos TMT-MS2 showed relative higher recovery of LFQ-DIA-Astral DE candidates, ∼79.3% of detection recovery, ∼75.7% of direction concordance, ∼60.9% of strict statistical support across the three pairwise genotype comparisons (Figure 3), suggesting a practical strategy for laboratories without access to Ascend or Astral instrumentation. In addition, we observed the lower cross-platform recovery of LFQ-DIA-Astral DE candidates in the comparison between *daf-2(lf);daf-16(lf)* and WT, which may reflect Astral DIA’s greater ability to quantify less accessible or subtler protein changes, as *daf-2(lf);daf-16(lf)* and WT are expected to be more similar proteomically.

Our results suggest that workflow selection should be guided by the experimental goal and available instrumentation rather than by identification depth alone. When Astral instrumentation is available, LFQ-DIA-Astral provides the broadest single-method LFQ coverage in this study. When Astral is unavailable but Orbitrap Ascend is available, the combination of LFQ-DIA-Ascend and TMT-MS2 provided the next-best alternative, recovering most of the candidate information obtained with LFQ-DIA-Astral; LFQ-DIA-Ascend alone offers the strongest single-platform approximation of Astral-derived candidates. For laboratories limited to Lumos workflows, combined Lumos DIA and Lumos TMT-MS2 analysis provides a practical alternative by integrating DIA coverage with TMT quantitative reproducibility. Because no workflow or workflow combination achieved complete Astral-candidate recovery, these alternatives should be viewed as fit-for-purpose strategies for recovering and prioritizing major protein changes, rather than full substitutes for LFQ-DIA-Astral depth. This comparison reflects relative recovery and concordance of candidates across methods, rather than an assessment against a ground-truth reference and should be interpreted as practical guidance for workflow selection rather than a benchmark of quantitative accuracy.

### 3.3 | Functional analysis of candidate proteins identified by comparative proteomics

We further performed functional categorization of the identified protein candidates, and found that, among the up-regulated proteins in *daf-2(lf)* compared to WT, the category related to extracellular material in collagen was enriched (Figure 4A). The STRING (version 12.0) (Szklarczyk et al., 2023) protein-protein interaction network further revealed that these collagen proteins form functional clusters (Figure 4B). This proteome-level finding is consistent with a previous study showing that collagen genes are transcriptionally up-regulated in the *daf-2(lf)* mutant and are required for its lifespan-extending effect (Ewald et al., 2015). Another functional group enriched in the up-regulated proteins was metabolism related to lipid and short chain dehydrogenase, which supports altered lipid metabolism in the long-lived *daf-2(lf)* mutant (McElwee et al., 2004; Mutlu et al., 2021). Specifically, the up-regulation of proteins K07B1.4 (monacylglycerol O-acyltransferase) and MBOA-2 (diacylglycerol O-acyltransferase) required for triglyceride biosynthesis may underline the increased fat storage in the *daf-2(lf)* mutant (Ashrafi et al., 2003). Furthermore, the up-regulation of ACBP-1 (fatty acyl-CoA binding protein), F07F6.7 (apolipoprotein L3), LBP-6 (fatty acid binding protein), LBP-7 (fatty acid binding protein), FAR-3 (fatty acid/retinol binding protein), NLT-1 (non-specific lipid transfer protein), ACOX-1.4 (acyl-CoA oxidase), and ACOX-1.5 (acyl-CoA oxidase) involved in lipid transport and oxidation indicates that the *daf-2(lf)* mutant may actively utilize lipids in its metabolic process (Figure 4C). In addition, we found that the stress response functional groups specifically involved in detoxification and oxidative stress are upregulated in *daf-2(lf)* compared to WT (Figure 4D and 4E), including cytochrome P450 family (CYPs), glutathione S-transferase (GSTs), and UDP-glucuronosyltransferases (UGTs). These proteins are key components of the cytoprotective mechanism, some of which contributes to the increased longevity in the *daf-2(lf)* mutant.(Shore & Ruvkun, 2013) Interestingly, previous studies also suggested that DAF-16 is required for the transcriptional up-regulation of some of these stress response genes (Dues et al., 2017; Huang et al., 2022; McElwee et al., 2004; Murphy et al., 2003).

**Figure 4.**
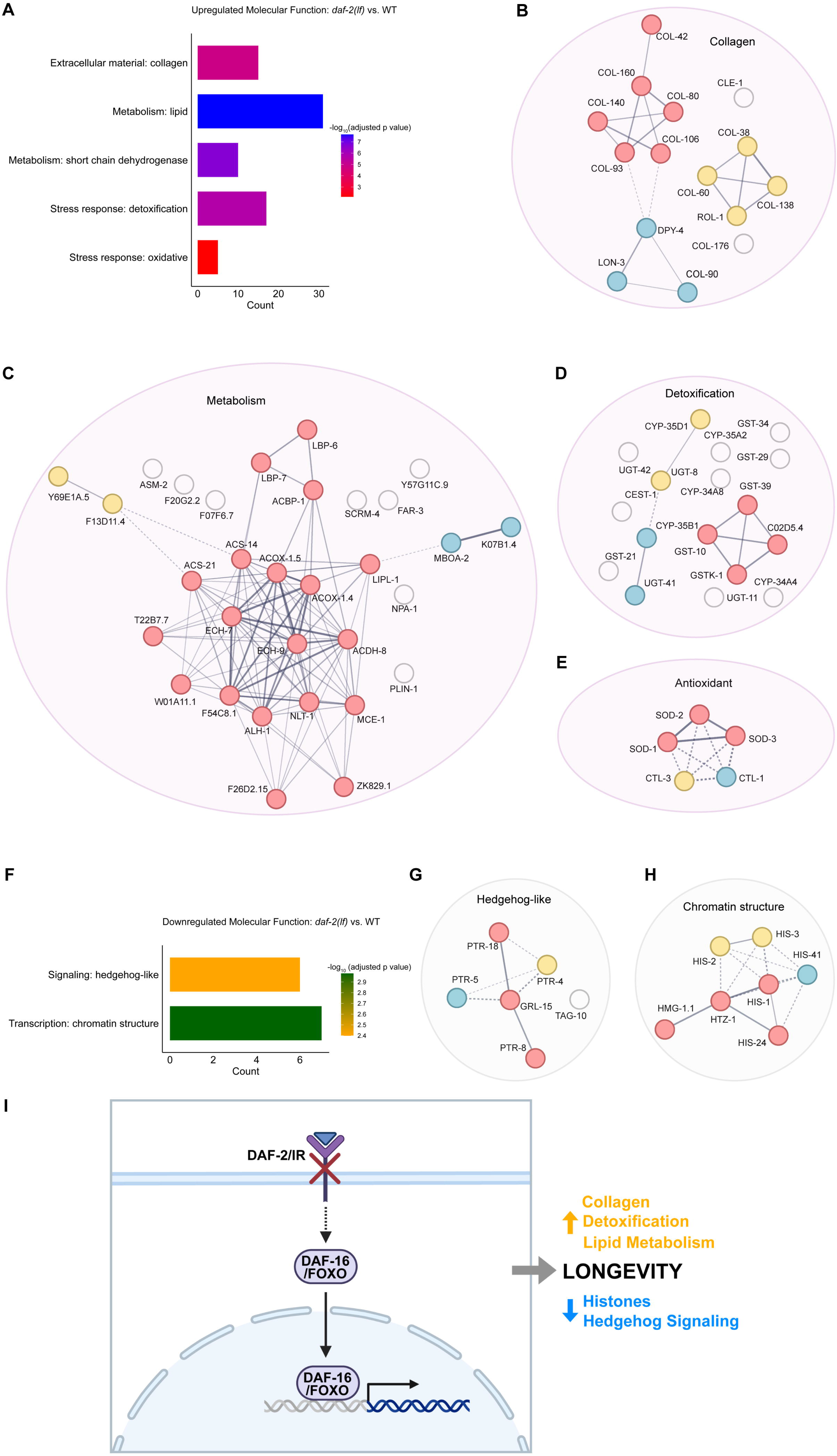
Functional mapping of differentially expressed proteins linked to IIS longevity. (A) Bar plot shows enriched biological processes among candidates upregulated in *daf-2(lf) vs*. WT. (B-E) STRING analysis of candidates up-regulated in *daf-2(lf) vs*. WT reveals distinct clusters of interacting proteins, with interconnected collagen family members (B), metabolic factors (C), detoxification-related enzymes (D) and antioxidant enzymes (E). (F) Bar plot shows enriched biological processes among candidates downregulated in *daf-2(lf) vs*. WT. (G, H) STRING analysis of candidates down-regulated in *daf-2(lf)* vs. WT reveals distinct clusters of interacting proteins enriched for hedgehog signaling (G) chromatin structure (H). (I) Summary model of proteomic changes associated with reduced insulin/IGF-1 signaling-mediated longevity. (STRING analysis: solid lines connect proteins within the same cluster; the thickness represents the combined confidence score of interaction between two proteins; dashed lines connect proteins of inter-cluster interaction. The threshold of confidence is set as medium. The k-means clustering with three clusters is applied.)

On the other hand, when analyzing the down-regulated proteins in the *daf-2(lf)* mutant, we observed enrichments in hedgehog-like signaling and transcription-chromatin structure (Figure 4F). PTRs are patched-like hedgehog receptors in *C. elegans*, and several of them form a cluster centered around GRL-15, a hedgehog related protein homolog, in the STRING network (Figure 4G). The hedgehog-like signaling has been associated with longevity regulation, and RNAi knockdown of *ptr-8* specifically in the germline or intestine, extends *C. elegans* lifespan.(Zhou et al., 2024) Thus, down-regulation of the protein components in hedgehog-like signaling may contribute to the lifespan extension in the *daf-2(lf)* mutant. In addition, chromatin structure changes are hallmarks of aging, including alterations in histone modifications and chromatin remodeling (Wang et al., 2022). We found that several histone proteins as well as HMG, the non-histone chromosomal protein are down-regulated in *daf-2(lf)* mutant compared to WT (Figure 4H), and form a cluster in the STRING network. These results indicate that the *daf-2(lf)* mutant may carry a global change in chromatin structure and histone modification landscape, specifically shaping the transcriptional program in this long-lived state.

### 3.4 | Screening for longevity promoters from proteomic dataset

Next, we focused on the protein candidates that were downregulated in *daf-2(lf)* compared to WT or *daf-2(lf);daf-16(lf)*, but not differentially changed between *daf-2(lf);daf-16(lf)* and WT using the criteria described above (Table S6 (Excel sheet 5)), searching for those whose inactivation promotes longevity. In total, 166 protein candidates were selected. Among them, 37 proteins have no human orthologs (Hu et al., 2011), and 36 proteins have already been reported to modulate lifespan or to be DAF-16 target genes (Table S8). We thus focused on the remaining 93 proteins and knocked down their encoding genes using RNAi and examined their effects on WT lifespan (Table S6 (Excel sheet 6)). Based on our RNAi screening results (Figure S8), we identified seven candidates and further validated their effects using longitudinal lifespan assays. We found that RNAi inactivation of *taf-2*, *clec-10*, and *F44E2.4* led to 22%, 15%, 14% of mean lifespan extension, respectively (Figure 5A-5C, Table S9), while RNAi inactivation of *cat-4*, *agr-1*, *K10D3.4* showed no lifespan changes (Figure 5D-5F, Table S9). TAF-2 is a transcription factor, a homolog of transcription initiation factor TFIID subunit 2 (TAF2), which plays a major role in the initiation of RNA polymerase II-dependent transcription.(Chen et al., 2021) It is known that TAF-2 acts with TAF-1 in broad embryonic transcription (Walker et al., 2004). One previous study shows that TAF-4, a member of TAF-family protein, is required for the life extension of *isp-1*, *clk-1* and *tpk-1* mitochondrial mutants, which are in the mitochondrial longevity context (Khan et al., 2013). A plausible interpretation is that *taf-2* knockdown may beneficially dampen broader transcriptional output after embryogenesis, reducing energetically costly and pro-aging gene expression without causing overt loss of fitness. However, its biological mechanism requires further investigation. CLEC-10 is a C-type lectin, which is predominantly expressed in the intestine based on the snRNA-seq analysis (https://agingc.shinyapps.io/AgingC_expression/). (Gao et al., 2024) Given the broader placement of C-type lectin genes in pathogen-response and immune-regulatory programs (Pees et al., 2021; Troemel et al., 2006; Youngman et al., 2011), the most plausible interpretation is that *clec-10* normally promotes a host-bacteria interactions state that is beneficial during infection but costly under standard laboratory conditions. It is reported that CLEC-10 is also downregulated in the long-lived *daf-10(m79)* mutant (Gaglia et al., 2012). F44E2.4 shows low homology to human low density lipoprotein receptor-related protein. In the previous RNA-seq transcriptomic analysis, the gene encoding CLEC-10 was transcriptionally down-regulated in the *daf-2(lf)* mutant compared to wild-type worms, while the F44E2.4 encoding gene was up-regulated, and the TAF-2 encoding gene showed no transcriptional changes (Zhang et al., 2022). Thus, some proteins relevant to *daf-2(lf)-*mediated longevity may exhibit significant changes in abundance at the protein level, even when their transcript levels remain unchanged or change in the opposite direction. Proteome-level analysis is therefore essential for uncovering longevity regulators that may be overlooked by transcriptomic analyses.

**Figure 5.**
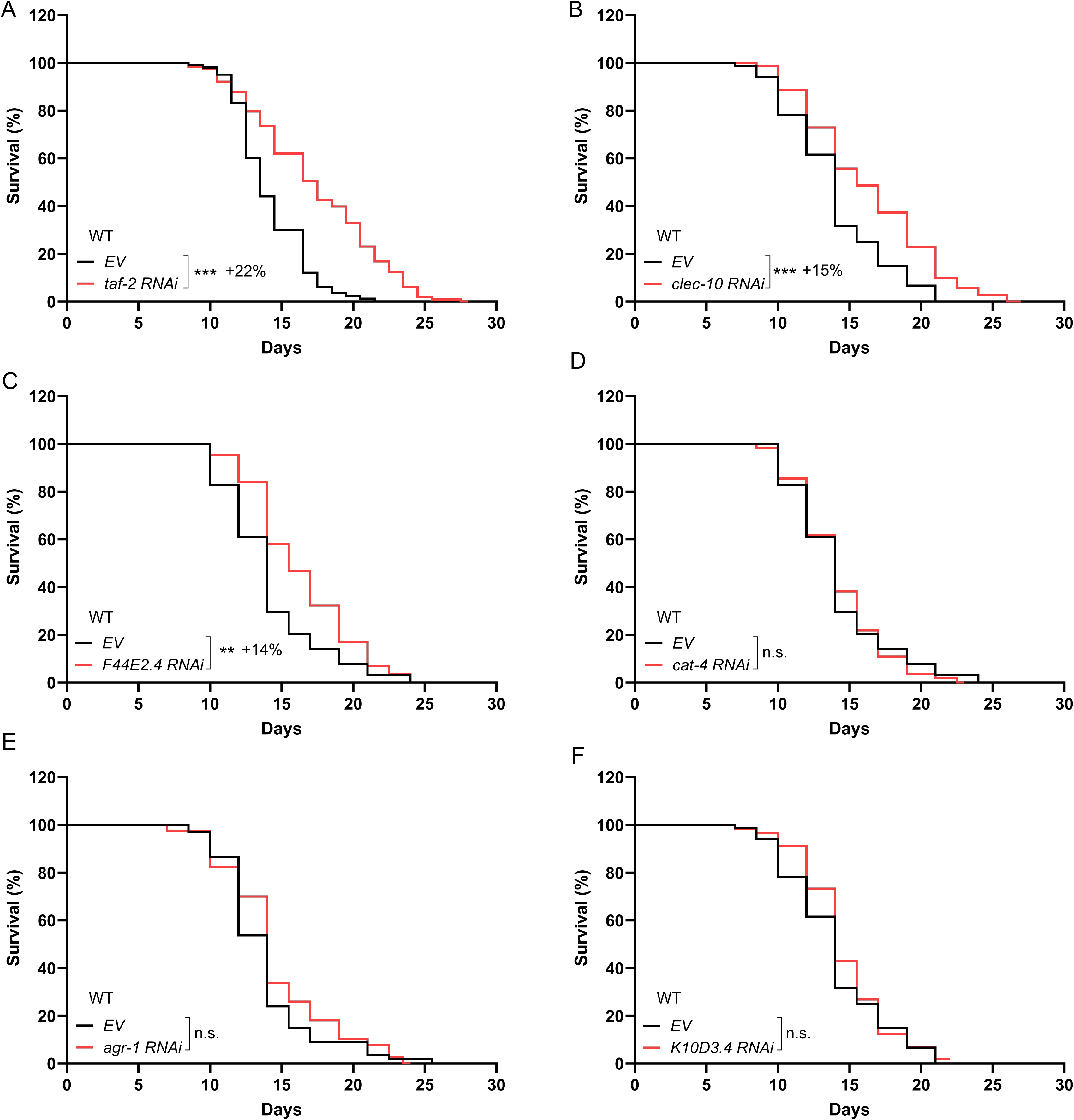
Lifespan assessment of protein candidates from genetic screens. RNAi inactivation of *taf-2* (A), *clec-10* (B), *F44E2.4* (C) extends mean lifespan by 22%, 15%, 14% respectively in WT worms, but RNAi inactivation of *cat-4* (D), *agr-1* (E), *K10D3.4* (F) does not affect WT lifespan. (n.s. = not significant p>0.05, *p≤0.05, **p≤0.01, ***p≤0.001, by log-rank test; n = 3 biological independent replicates for A-C, n = 2 biological independent replicates for D-F, 60∼120 worms per replicate, see Supplementary Table S9 for full lifespan data.).

## 4 | DISCUSSION

MS–based proteomics has become a routine and powerful approach for advancing biological and biomedical research, driven by rapid advances in both instrumentation and bioinformatics. Previous studies have systematically benchmarked quantitative proteomics and phosphoproteomics workflows. Our study builds on this foundation but addresses a different question how these workflows perform in a biologically complex, whole-organism longevity model and how method choice influences candidate prioritization. By evaluating three common methods—TMT, LFQ, and SILAC-spike-in, alongside different mass spectrometers (Lumos, Ascend, and Astral) used for LFQ-DIA, we found that many candidates showed consistent directions of change across workflows, which supports the use of cross-method trend consistency validation. While each approach and instrument combination showed unique strengths and limitations regarding sensitivity, throughput, quantification accuracy, and proteome coverage.

For TMT, although the TMT-SPS-MS3 workflow mitigates ratio compression relative to TMT-MS2, we still recommend using relaxed fold-change cutoffs to account for residual ratio compression. Notably, the TMT reporter ion isotopic correction factors were not applied in the MSFragger analysis within FragPipe, as the platform currently lacks this capability. LFQ-DDA often suffers from poor reproducibility for low-abundance proteins. SILAC and its variants require specialized conditions to label amino acids such as arginine and lysine, and a biological system with significant arginine-to-proline conversion can perturb the analysis. The growth difference in different label conditions is informative when working on organisms whose development could be sensitive to heavy labeling. Moreover, multiplexed labeling demands careful forward and reverse labeling schemes and high labeling efficiency, or an alternative SILAC-spike-in strategy used in this study. In addition, the SILAC method yielded the lowest proteome coverage. Although incorporating extensive fractionation can mitigate this limitation, it substantially increases instrument run time and requires a larger amount of starting materials. LFQ-DIA has advanced rapidly due to improved software pipelines, but the choice of instrument can dramatically influence peptide identification depth and data quality. Although we optimized liquid chromatography and MS parameters for the Ascend instrument, resulting in nearly 8,000 identifications, the wider m/z window and longer LC gradients differ considerably from the Astral setup. We adopted a library-free strategy for LFQ-DIA, yet a robust, library-based approach could improve accuracy at the expense of proteome coverage. For LFQ-DIA optimization on Lumos, we found that when we applied tMS2 mode rather than regular DIA mode with the desired isolation window provided more identified proteins. However, the intrinsic difference and influence for quantification between these two modes with the same isolation window require further investigation.

In addition to the general features of LFQ-DIA among different instruments, the instrument itself has different strengths and weaknesses. For example, the Fusion Lumos is a well-proven multi-purpose instrument with relative lower cost (older version), supporting advanced fragmentation and MS^n^ while slower acquisition and lower throughput; the Ascend has high speed and sensitivity and maintains Orbitrap multianalysis versatility but slightly more complex with relatively higher cost (relatively new version); the Astral has extreme throughput and sensitivity but only supports MS1 and MS2 and does not support MS^n^ with very high cost (the newest version).

In general, the choice of proteomic method and instrument must be carefully aligned with the specific experimental goals, such as maximum coverage, reproducibility, or accurate quantitation. Ultimately, our findings reinforce the idea that no single technique can comprehensively address all questions in complex systems. Adopting a comparative framework not only enhances confidence in shared outcomes but also allows the discovery of novels that might go unnoticed with a single-method approach. Even though our study focused on quantitative proteomic methods comparison, and many of the proteins and pathways discussed above have been previously implicated in longevity regulation, we identified three proteins that were not previously linked to longevity regulation. Molecular mechanisms underlying their effects are worth further investigation.

## AUTHOR CONTRIBUTIONS

L.D. and M.C.W. conceptualization; L.D. data curation, L.D. formal analysis; M.C.W. funding acquisition; L.D., H.L., C.H., V.V., K.A., S.R.P. investigation; L.D., H.L., A.S.M, Y.Z., X.M., J.M.C., F,J. methodology; J.W. and M.C.W. resources; L.D. software; M.C.W. supervision; L.D. and M.C.W. validation and visualization; L.D. writing-original draft; L.D. and M.C.W. writing-review and editing.

## ACKNOWLEDGEMENTS

M.C.W. was supported by the Howard Hughes Medical Institute (HHMI). L.D. was supported by the HHMI Janelia Visiting Scientist Program. We are grateful to Dr. He Zhu for valuable suggestions, and to Dr. Sung Yun Jung for kind technical guidance. We appreciate Dr. Nan Wang for maintaining the Orbitrap Fusion Lumos and the Orbitrap Ascend Mass Spectrometers when the instruments were at Janelia Research Campus, HHMI. We thank Laura Hart, Princeton Svay, Olivia DeThomasis, Hina Naz, Brian Rebhorn and Amanda Cavallaro for maintenance support; BioRender for the support on creating Figures 1A, 1F, and 1O; and the Caenorhabditis Genetics Center (CGC) for C. elegans and bacteria strains, as well as WormBase.

## DATA AVAILABILITY STATEMENT

All the LC–MS raw files have been deposited into MassIVE (ftp://MSV000098193@massive-ftp.ucsd.edu. Password: CP2025). Any additional information required to reanalyze the data reported in this paper is available from the lead contact upon request.

